# Phage peptide promotes bacterial cell death via inner and outer membrane targets

**DOI:** 10.64898/2026.11.12.765902

**Authors:** Michael A. Debrah, Luis A. Carrillo, Binaya K. Baral, Makahla Schnarre, Luke Wardrip, Gary A. Lorigan, Andra B. Schromm, Jolene Ramsey

**Affiliations:** Department of Biology, Texas A&M University, College Station, Texas, 77845, USA; Center for Phage Technology, Texas A&M University, College Station, Texas, 77845, USA; Department of Biochemistry, Texas A&M University, College Station, Texas, 77845, USA; Department of Chemistry and Biochemistry, Miami University, Oxford, Ohio, 45056, USA; Division of Immunobiophysics, Priority Area Infections, Research Center Borstel, Leibniz Lung Center, 23845 Borstel, Germany

**Keywords:** antimicrobial peptide, cationic, membrane, lysis, bacteriophage, bacteria, disruptin

## Abstract

Cationic antimicrobial peptides (CAMPs) play a major role in mammalian innate immunity by preferentially targeting bacterial membranes through charge-dependent interactions. While CAMPs are extensively studied in eukaryotic systems, less is known about phage-encoded CAMPs and their role in bacterial lysis. Here, we characterize gp28, a 56-aa cationic peptide encoded by the T7-like coliphage phiKT to define its mechanism of membrane disruption and bacteriostatic activity. Biochemical fractionation reveals that gp28 is asymmetrically distributed across bacterial membranes. Supporting its direct role in causing cell death, gp28 induces permeability changes at the inner membrane. Using synthetic gp28 peptide, we also show that gp28 activity is blocked by lipopolysaccharide and divalent cation competition. This led us to discover that gp28 is also active against Gram-positive bacteria in a surface charge-dependent manner. Finally, we demonstrate gp28 has broad bacteriostatic activity against multiple Gram-negative species, including clinical Uropathogenic *E. coli* (UPEC) isolates. Together, our results support a model in which gp28 exploits charge-based interactions with bacterial surface molecules to gain access to membranes, then destabilizes the envelope to promote host cell death. These findings expand our understanding of phage lysis strategies and highlight phage-derived peptides as potential scaffolds for antimicrobial development.

**SIGNIFICANCE STATEMENT:** Cationic antimicrobial peptides produced by eukaryotes target and disrupt bacterial membranes. These peptides are integral members of the innate immune system and potential antibacterials. Due to their cellular origin, the genetics to understand their mechanisms of action are cumbersome. Here, we demonstrate functional similarity between a bacteriophage peptide called gp28 and cationic antimicrobial peptides. Gp28 partitions between both membranes of Gram-negative bacteria and reduces their integrity. We show that gp28 employs charge-based interactions to associate with and disrupt membranes, suggesting that gp28 targets charged polymers on the membrane surface. Additionally, gp28 inhibits the growth of Gram-negative and Gram-positive bacteria, and renders them more susceptible to antibiotics. Our studies provide a foundation for future genetics and development of phage peptide-based antimicrobials.

## Introduction

Cationic Antimicrobial Peptides (CAMPs) are short, positively charged peptides produced in many mammals. Within an organism, CAMPs regulate inflammation and other innate immune responses(1). At the cellular level, they form a crucial component of the innate defense against bacterial infections. CAMPs typically range from 12-50 amino acids and carry a net positive charge that promotes preferential binding to the negatively charged surface of bacteria over the largely zwitterionic membrane of eukaryotic cells(2, 3). After initial binding, CAMPs disrupt bacterial membranes.

Two membrane-disrupting CAMPs used clinically to treat Gram-negative bacteria are Polymyxin E (colistin) and Polymyxin B (PMB). The polymyxins are comprised of a cationic heptacyclic peptide component, coupled to an extracyclic peptide chain with an acyl tail. Their acyl tail facilitates membrane insertion and the cationic component binds lipopolysaccharides (LPS) with high-affinity, displacing divalent cations such that outer membrane (OM) integrity is reduced. Similarly, ribosomally synthesized peptides, including human cathelicidin LL-37, contribute to host defense by targeting bacterial envelopes through similar charge-dependent interactions(4, 5). While most work on CAMPs has focused on cell-derived peptides, phages also use cationic, membrane permeabilizing peptides in bacterial lysis to release progeny virions.

Bacteriophages at the end of their lytic cycle must overcome a multilayered bacterial envelope to release progeny virions. Gram-negative bacteria have three layers of their envelope: the inner membrane, the peptidoglycan layer, and the outer membrane. The canonical lysis genes of double-stranded DNA phages for targeting Gram-negative hosts are holins, responsible for initiating lysis by making holes in the inner membrane (IM), endolysins for degrading the peptidoglycan layer, and spanins for disrupting the OM(6). Holins are small IM proteins that eventually disrupt the proton motive force (PMF)(7–9). To do so, holins form micron-scale holes after reaching a critical concentration to allow for endolysin entry into the periplasmic space(10). Endolysins are muralytic enzymes that degrade cell wall PG. There are two major types of endolysins: soluble and signal-arrest release (SAR) endolysin. Soluble PG hydrolases accumulate in their active state in the cytoplasm until holin pores give them access to their periplasmic substrate(6, 10). In contrast, SAR endolysins are simultaneously released into the periplasmic space after membrane polarization, where they assume an active conformation(11). Finally, an active OM disruption step is essential to release the phages. One strategy uses spanin proteins, which are thought to induce inner and outer membrane fusion. Another strategy we recently identified in the phage phiKT is the use of a CAMP(12).

PhiKT is a T7-like dsDNA tailed phage that infects *E. coli* with modified O22-LPS, such as found in the 4s strain(13). Sandwiched between the holin and endolysin genes of PhiKT is the CAMP gene, called gp28 or the disruptin(12). Gp28, a 56-amino acid peptide, is primarily alpha-helical with a net positive charge of +8 at physiological pH(12, 14, 15). We previously demonstrated that gp28 is membrane localized and complements the lysis defect of a spanin-deficient lambda lysogen(12). Also, when synthetic peptides were added externally in bactericidal and inhibitory experiments against the Gram-negative *E. coli* strains, gp28 was active at concentrations equivalent to the mammalian LL-37(12). These observations suggest that gp28 functions as a phage-encoded CAMP that can disrupt the load-bearing OM barrier and cause cell death independent of holin and endolysin activity. However, the mechanism by which it disrupts the OM, and the role of its cationic properties remain poorly understood.

In this report, we demonstrate that gp28 charge-based interactions with the membrane are key to its membrane disruption properties. We show that gp28 is enriched in the IM, disturbs IM integrity, and induces permeability changes that are sufficient to trigger early holin holes. Using synthetic peptide, we demonstrate the bactericidal activity of gp28 against a panel of Gram-negative bacteria, clinical Uropathogenic *E. coli* (UPEC) strains, and Gram-positive bacteria with varied surface charge.

## Results

### Gp28 is present in both membranes with predominance in the IM

We previously showed using immunofluorescence that gp28 has an IM localization pattern(12). However, we also demonstrated that gp28 complements the OM disruption defect of a spanin-null lambda lysogen. If gp28 has a direct effect on the OM, we hypothesized that gp28 would localize in the OM during phage lysis. We measured gp28 abundance in sucrose-gradient fractionated bacterial membranes (Fig. 1A). Gp28 was distributed ∼80% with the IM fraction and ∼20% with the OM fraction (Fig. 1B-D).

**Figure 1.**
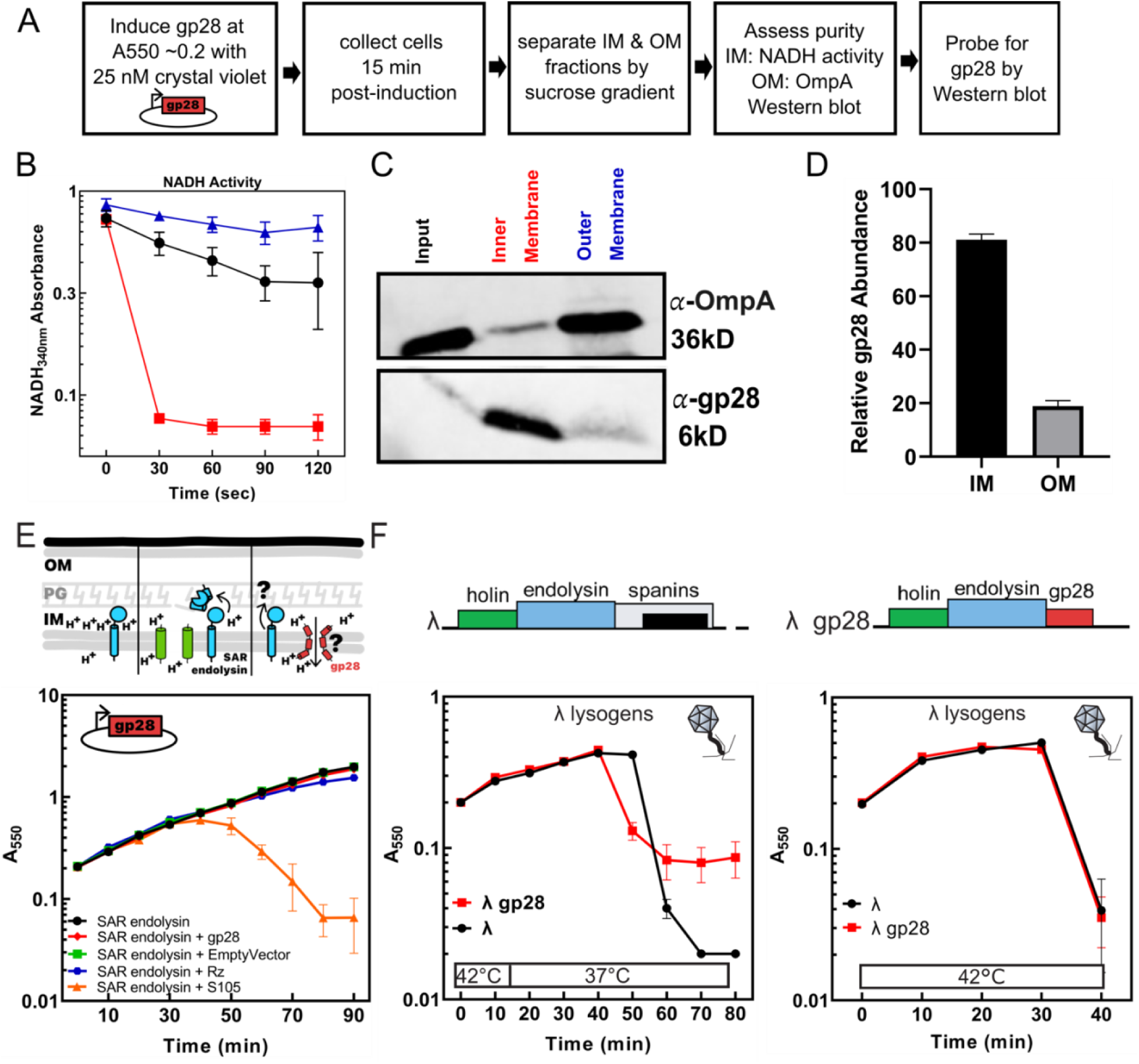
Gp28 primarily localizes to the IM. **(A)** Membrane fractionation workflow for plasmid-based gp28 expression. Isolated membrane fractions were assayed for **(B)** IM NADH dehydrogenase activity and **(C)** OM OmpA or gp28 abundance by western blot. **(D)** Densitometry quantification of gp28 band intensities from 3 biological replicates. Averages were normalized to the total gp28 detected per experiment and plotted with Standard Error of the Mean. **(E)** Schematic of the SAR endolysin in the IM. *E. coli* MG1655 carrying a pBAD33 gp22 under a P_BAD_ promoter expressing the SAR endolysin alone (black circle) or with pBAD24 empty vector (green square), pBAD24 pS105 (orange triangle), pBAD24 gp28 (red diamond), pR’Rz + pQ-Kan (blue hexagon) induced by 0.2% arabinose. **(F)** Schematics of lysis cassette in lambda and lambda gp28 lysogens. *E. coli* MG1655 lambda (black circle) and lambda gp28 (red square) lysogens were induced as indicated on the graph and monitored for lysis. The average of three biological replicates were plotted with the Standard Error of the Mean.

These data are consistent with the functional roles seen for gp28 across the membranes. The previously observed activity of gp28 in the IM was increased permeability to small DNA dyes(12). One explanation for increased dye permeability is pore formation. We used the detection of SAR endolysin activity as a sensitive assay for detecting IM pores. When pores ablate the PMF, the SAR domain is released, and PG is degraded (Fig. 1E). Co-expression of SAR endolysin with the S105 holin, but not a nonspecific IM protein the spanin Rz, results in cell lysis (Fig. 1E). In contrast, co-expression of SAR endolysin with gp28 did not result in cell lysis, suggesting that gp28 alone does not form pores Fig. 1E).

An alternative explanation for the gp28-mediated change in membrane permeability is a temporary perturbation in the electrochemical gradient driving the PMF. A sensitive assay for detecting small changes in the electrochemical gradient across the inner membrane is the timing of holin hole formation. Holin triggering is genetically programmed but can also be induced artificially by protonophores that dissipate the PMF. To test whether gp28 interrupts the PMF, we replaced the spanins in a lambda (λ) prophage with gp28 and assayed lysis timing. Under the standard thermal induction conditions, gp28 caused early lysis, suggesting that the permeability changes induced by gp28 are sufficient to trigger early holin holes (Fig. 1F). Interestingly, when cells were maintained at higher induction temperatures throughout the experiment, which does not permit as much gp28 accumulation prior to holin triggering, the lysis timing was not accelerated. Overall, these results support a model where the gp28 in the IM disrupts its integrity and affects phage lysis.

### Gp28 activates an OM stress response pathway

The gp28 that localizes to the OM is destabilizing its load-bearing properties. A major structural component contributing to the OM mechanical stability are LPS-LPS salt bridges with divalent cations (16–18). Displacement of divalent cations from negatively charged LPS by cationic peptides, such as PMB, is known to destabilize the OM and trigger the Rcs (regulation of capsular polysaccharide synthesis) cell surface stress response pathway(19, 20). Due to the cationic nature of gp28, we hypothesized that it would disrupt LPS-LPS interactions and activate the RcsF sensor to initiate the Rcs phosphorelay. To test this, we used a luciferase-based transcriptional reporter for P*rprA*, an Rcs-responsive promoter (Fig. 2A). PhiKT infection producing native gp28 levels demonstrated low level activation of the Rcs stress response pathway coincident with lysis initiation (Fig. 2B). Interestingly, gp28 overexpression alone from a plasmid at sublethal and lethal levels did not activate the Rcs response (Fig. 2C). However, synthetic gp28 peptide resulted in dose-dependent increases in luciferase activity, albeit at lower levels than PMB-induced luciferase activity (Fig. 2D). Overall, these results suggest that gp28 in the OM disrupts surface LPS-LPS interactions to reduce its structural integrity during lysis.

**Figure 2.**
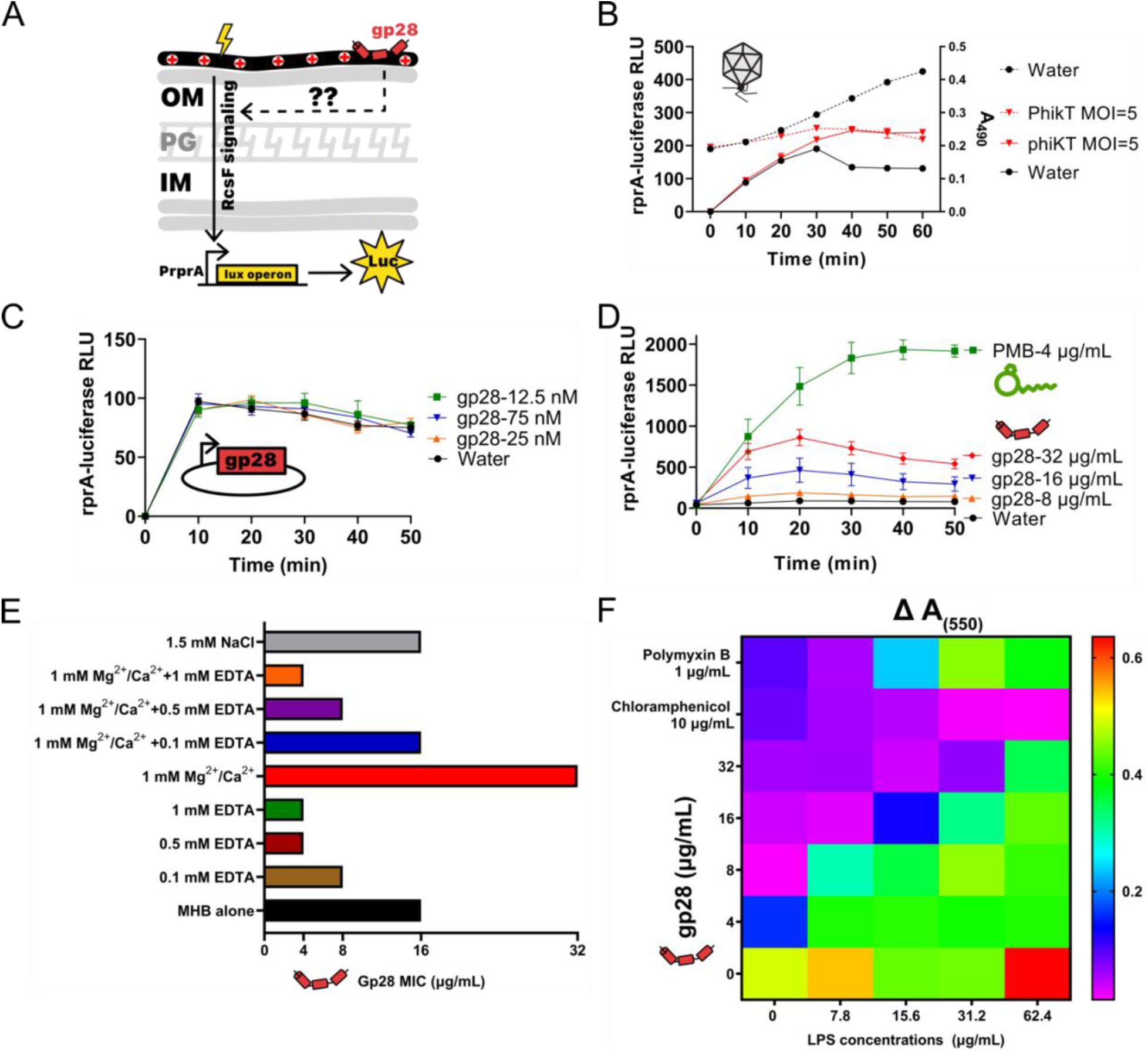
Gp28 triggers the Rcs OM stress pathway. **(A)** MG1655 carrying the pJW15 PrprA-lux plasmid with the native RcsF-responsive rprA promoter transcriptionally fused to a luciferase reporter were exposed to either **(B)** peptide expressed natively during phage infection, **(C)** plasmid-expressed peptide, or **(D)** externally added peptides and assessed for luciferase. **(B)** For external addition, cells were treated with water (black circle), gp28 at 8 ug/ml (orange triangle), 16 ug/ml (blue inverted triangle), 32 ug/ml (red diamond), or polymyxin B at 4 ug/ml (green square). **(C)** For plasmid expression, psJExD-spec-gp28 in which gp28 is expressed from the P_JExD promoter, were induced with crystal violet at 0 nM (black circles), 12.5 nM (green squares), 25 nM (orange triangles), or 75 nM (blue triangles). **(D)** Cells were infected with phiKT at an MOI of 5 (red inverted triangles) or treated with water as a negative control (black circles). Solid lines show PrprA-luciferase activity (RLU; left y-axis), whereas dashed lines show culture absorbance A_490_ over time. The Relative Luminescence Unit measurements representing at least 3 biological replicates is shown. **(E)** MIC for *E. coli* 4s cells grown in Mueller-Hinton Broth supplemented with divalent cation MgCl_2_ ^2+^ and CaCl_2_^2+^ salts, chelator ethylenediaminetetraacetic acid (EDTA), or monovalent NaCl salts for 16 h in the presence of synthetic gp28. The horizontal bars represent the consensus of the MIC after 3 biological replicates. **(F)** A heatmap of change in optical density for *E. coli* 4s grown with gp28, chloramphenicol, or PMB in the presence or absence of increasing concentrations of LPS in a standard 16h MIC assay. The average of three biological replicates were plotted.

### Gp28 antimicrobial activity is blocked by electrostatic competitors

Increasing the concentration of divalent cations strengthens the OM barrier and can reduce the activity of CAMPs by enhancing charge screening at the surface (21). We further explored the hypothesis that gp28 specifically interacts with LPS in the OM using synthetic peptide MIC assays. First, we saw that cation chelation by EDTA reduced the MIC by as much as 4-fold (Fig. 2E). Conversely, a higher concentration of the divalent cations Mg^2+^ and Ca^2+^ increased the MIC of gp28 from 16 to 32 ug/mL against *E. coli* 4s. Furthermore, both chelation and excess cations mimicked the chelator alone pattern. Consistent with these data, monovalent Na^+^ treatment did not affect the MIC. These data suggest that electrostatic interactions between gp28 and LPS on the cell surface are critical for its antimicrobial activity.

To more directly test the hypothesis that gp28 would disrupt LPS-LPS interactions in the OM, we employed an LPS blocking assay. The blocking assay added LPS isolated from *E. coli* 4s as a competitor in an MIC assay. Chloramphenicol, a neutral antibiotic, was unaffected by added LPS (Fig. 2F). However, both PMB and gp28 MICs against *E. coli* 4s, the native phage phiKT host, were reduced by LPS in a concentration-dependent fashion (Fig. 2F). These data suggest excess LPS in the supernatant may directly inhibit gp28 access to the membrane, consistent with a charge-based interaction between LPS in the OM and gp28.

### Gp28 increases cell susceptibility to antibiotics

Prior work with small, cationic dyes suggested that gp28 increases permeability across both the IM and the OM(12). We predicted that gp28 would also increase cell susceptibility to antibiotics, including those otherwise excluded by the outer membrane due to size. To test this prediction, we expressed a nonlethal level of intracellular gp28 that confers outer membrane disruption by our complementation assay and measured changes in disk assay zone of inhibition for a broad panel of antibiotics. Across the five classes of antibiotics tested, most displayed an increased zone of inhibition in the presence of gp28 (Fig. 3A). Although the β-lactam derivatives: aztreonam, meropenem, cefepime, and ceftazidime, and other targeted antibiotics sulfamethoxazole, nalidixic acid, and polymyxin B had increased zones of inhibition, only the doxycycline increase reached statistical significance (Fig. 3A). These data are consistent with a model whereby low, nonlethal levels of gp28 permeabilize *E. coli* membranes.

**Figure 3.**
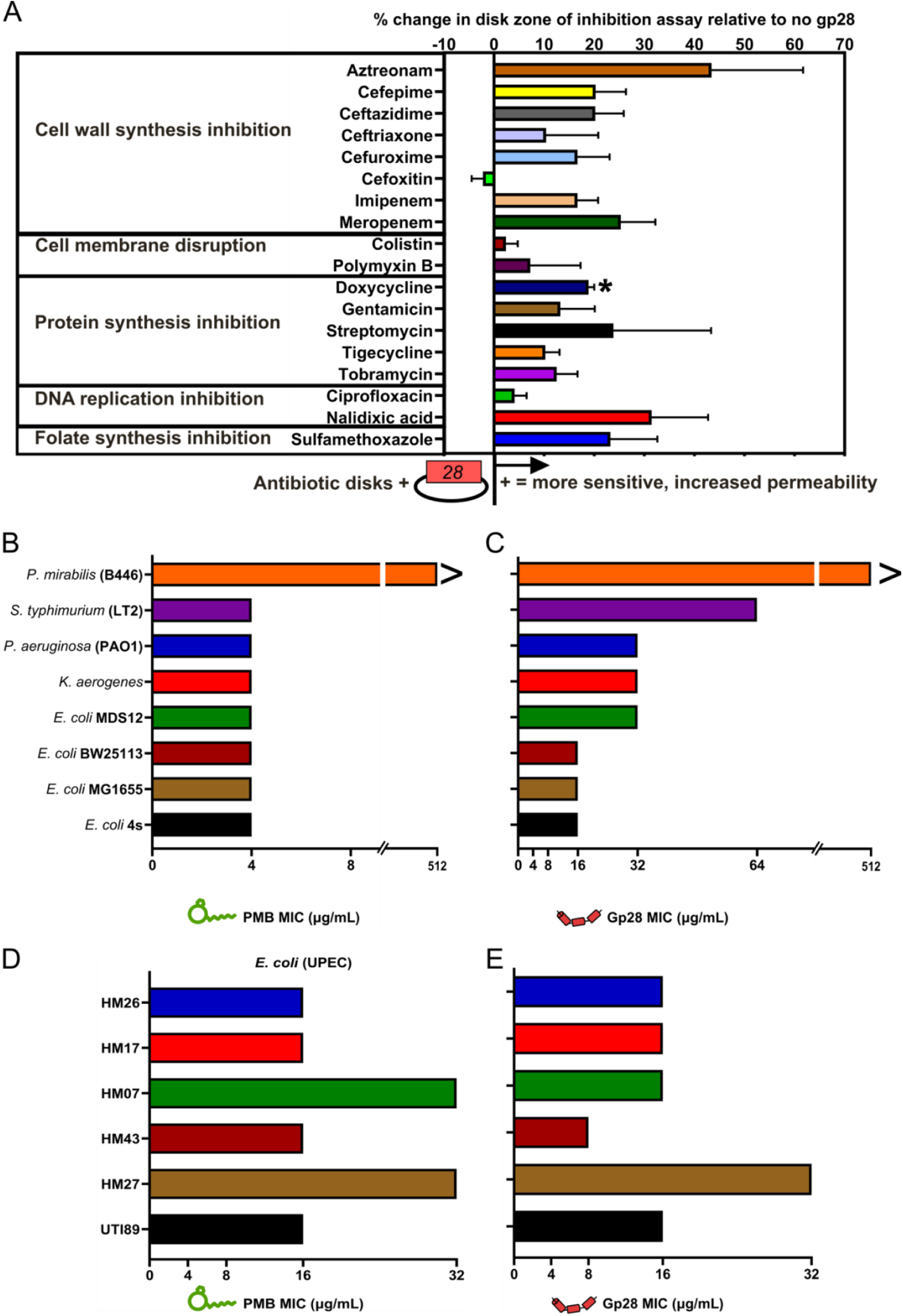
Gp28 inhibits growth of multiple Gammaproteobacterial pathogens. **(A)** *E. coli* MG1655 carrying pBAD24 empty vector or including a gp28 insert were plated in lawns and induced with 0.2% arabinose. Antibiotic disks were placed on lawns prior to incubation (full details in methods). The graph is plotted as the difference of the measured pBAD24 gp28 zone of inhibition from that of pBAD24. A two-sided one-sample t-test was performed. A p-value below 0.05 is marked by an *. **(B-E)** MIC for the indicated panel of Gram-negative species and selected Uropathogenic *E. coli* (UPEC) isolates in the presence of either PMB **(B, D)** or gp28 **(C, E)** peptide. Horizontal bars indicate the consensus MICs derived from three independent biological replicates. The X axes of panels A and B are truncated before 512 for clarity.

### Gp28 inhibits growth across the *Gammaproteobacteria*

Cationic peptides are broadly active across Gram-negative bacteria. Previously, we reported that gp28 was bactericidal against both the MG1655 *E. coli* lab strain and the 4s equine isolate. Here, we extend that finding with the standard MIC assay testing additional *E. coli* strains and four other Gram-negative bacteria. All the tested strains except *Proteus mirabilis* were susceptible to PMB at an MIC of <4 ug/mL (Fig. 3B). *E. coli* MDS12, a strain with all active prophage regions deleted, had an MIC of 32 ug/mL, higher than the 16 ug/mL in MG1655, BW25113, and 4s (Fig. 3C). *K. aerogenes* and *P. aeruginosa* growth was also inhibited at 32 ug/mL, while *S. typhimurium* was not inhibited until 128 ug/mL of gp28. However, gp28 was not bacteriostatic at the maximum concentration of 512 ug/mL against *P. mirabilis* (Fig. 3B&C). We then assayed gp28 against six Uropathogenic *E. coli* (UPEC) clinical isolates using an MIC assay. Each of the UPECs tested were susceptible to 16-32 ug/ml PMB (Fig. 3D). The MIC of gp28 for all the UPEC clinical isolates was 16 ug/mL, except in *E. coli* HM43 where the MIC was 8 ug/mL (Fig. 3E). These data demonstrate that gp28 and PMB are active against the same bacteria, although the gp28 MIC is higher in all cases, demonstrating that gp28 is not selective for *E. coli* membranes.

### Gp28 is lethal to cells in a surface charge-dependent manner

We reasoned that if gp28 was not selective for *E. coli* membranes, it would integrate into any accessible membrane with a net negative surface charge. Since the wall teichoic acid and lipoteichoic acids of Gram-positive bacteria contain negatively charged phosphate groups (Fig. 4A), we hypothesized that gp28 would also be lethal to Gram-positive cells. We found that against *Bacillus subtilis* 168, the MIC of the CAMP Nisin is 4 ug/mL and the MIC of gp28 is 16 ug/mL (Fig. 4A). The *B. subtilis* net surface charge due to the wall and lipoteichoic acids is reduced with D-alanylation added by the gene products of the *dlt* operon(22–24). Four *B. subtilis dlt* gene knockout strain MICs were measured. For Nisin, the Δ*dltC* and Δ*dltD* strains showed the lowest MICs at 2 ug/mL, followed by 3 ug/mL for Δ*dltB*, and 3.5 ug/mL for Δ*dltA* compared to the parent MIC of 4 ug/mL for Nisin (Fig. 4A)(25). In contrast, after gp28 treatment *B. subtilis dlt* knockouts showed a fourfold lower MIC, reducing from 16 ug/ml to 4 ug/mL (Fig. 4A). All *B. subtilis* parent and mutant strains were susceptible to chloramphenicol, with an MIC of 2 ug/mL (Fig. 4A). These results demonstrate that gp28 lethal activity against Gram-positive cells is surface charge dependent.

**Figure 4.**
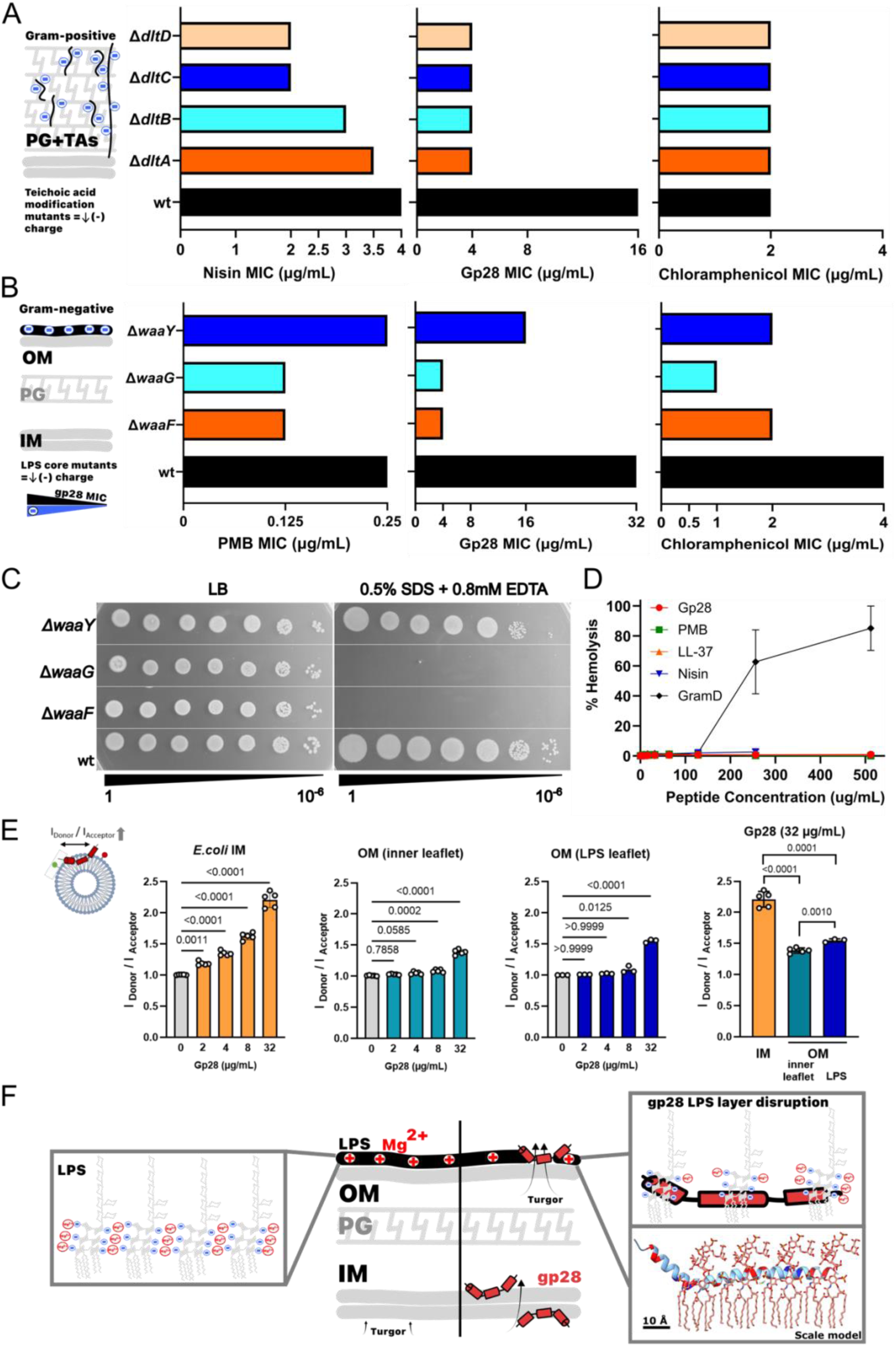
Gp28 is lethal to Gram-positive cells in a charge-dependent manner. **(A)** Schematic for Gram-positive *B. subtilis* surface charge distribution. *B. subtilis* 168 wt and Δ*dltA-D* mutant strains were subjected to the MIC assay in the presence of either Nisin, chloramphenicol, or gp28. **(B)** Schematic for Gram-negative *E. coli* surface charge distribution and the MIC assay for *E. coli* MG1655 wt, *ΔwaaF*, *ΔwaaG*, and *ΔwaaY*. **(C)** Tenfold serial dilutions of strains in (B) were spotted onto LB supplemented with 0.5% SDS and 0.8 mM EDTA, where indicated. **(D)** Bovine RBCs resuspended in PBS to a 2.5% final hematocrit were incubated with cationic antimicrobial peptides (gp28, PMB, LL-37, Nisin, Gramicidin D) at concentrations of 4, 8, 16, 32, 64, 128, 256, 512 ug/mL at 37°C for 60 min. The release of hemoglobin into the supernatant from RBC lysis was measured at 405 nm. Percent hemolysis induced by each peptide normalized to the PBS control. **(E)** Liposomal model membranes of the *E.coli* inner membrane (PE:PG:CL 81:17:2 M), the inner leaflet of the outer membrane (PE:PC 70:30 M), and the outer leaflet of the outer membrane (*E.coli* Ra LPS) were analyzed for gp28 membrane integration by Förster-resonance-energy-transfer (FRET) probe-dilution assay. Gp28 was titrated to the liposomes at 2, 4, 8, and 32 µg/mL at 37°C to liposomes in a cuvette under constant stirring. FRET ratios (I_Donor_/I_Acceptor_) are mean ±SD of n = 5 (IM, OM inner leaflet) and n = 3 (LPS) independent measurements. Data were analyzed by one-way ANOVA with Bonferroni post-hoc test. Comparison of end points were analyzed by unpaired two-sided Student’s *t*-test. **(F)** Model for gp28 destabilization of the Gram-negative envelope. Under physiological conditions, the turgor pressure generated by the cell interior is borne by the intact inner membrane, peptidoglycan, and outer membrane load-bearing layers. The OM barrier is stabilized by salt bridges between divalent cations and adjacent LPS molecules. gp28 localization to the membranes may displace cations in the LPS layer, compromising its load-bearing capacity and eventually leading to cell death. A scaled Alphafold3 model of gp28 with positively charged residues colored in red and the negatively charged residues in dark blue is shown manually positioned next to a chemical model of the LPS structure (Ra form). A single Mg2+ ion is shown (green).

We performed a parallel experiment in *E. coli*. The surface charge of the Gram-negative cell surface is reduced upon defects in LPS inner core biosynthesis, where both the first and second heptose are phosphorylated. We performed MICs in *E. coli* with defects in Δ*waaF*, which adds the second heptose moiety to the inner core, Δ*waaY*, the kinase that phosphorylates the second heptose in the LPS inner core saccharide, and Δ*waaG*, which initiates outer core synthesis and presents an 80% reduction in phosphorylation on the heptoses in the inner core (26). All three strains decreased their MIC by at least half for chloramphenicol, and by half for the PMB MIC in Δ*waaG* and Δ*waaF* (Fig. 4B). The Δ*waaG* and Δ*waaF* alleles were also sensitive to SDS/EDTA treatment (Fig. 4C), consistent with previous reports of compromised OM barrier functions in these mutants (27). The MIC for gp28 was reduced by half for ΔwaaY, but reduced 8x for Δ*waaG* and Δ*waaF* (Fig. 4B). Although the trend of the reductions in MIC is consistent with the B. subtilis result in Fig. 4A, the differences did not reach statistical significance. Employing liposomal model membranes, we analyzed gp28 interaction with the inner and outer membrane lipids by FRET-probing of the lipid bilayer. Titration of gp28 into liposomes demonstrated significant intercalation of gp28 into IM model membrane compositions at 2 µg/mL, with concentration-dependent membrane disturbance. In contrast, the OM inner leaflet and LPS model membranes required gp28 at MIC concentration (32 µg/mL) for intercalation, demonstrating significant higher attraction of gp28 to the IM and membrane disturbance below MIC as intrinsic properties of the bacterial lipid membranes (Fig. 4D).

Finally, we hypothesized that gp28 would not be active against eukaryotic cell membranes. Eukaryotic cell membrane lipids have an overall zwitterionic character, but the negatively-charged glycocalyx layer can attract CAMPs. To test this, we performed a hemolysis assay with bovine red blood cells (RBCs). PBS-treated RBCs remained intact, whereas treatment with 0.1% Triton X-100 caused complete hemolysis (Fig. 4E). Gramicidin D caused a dose-dependent hemolysis response, but PMB, Nisin, the mammalian LL-37, and gp28, like the PBS negative control, produced no hemolysis. Therefore, we conclude that gp28 does not cause lysis in bovine RBCs. Overall, gp28 is most active in bacterial cells with negatively charged surfaces.

## Discussion

In this study, we demonstrate that gp28 is localized to and functional in both the IM and OM. When exposed to gp28 synthetic peptide, cells respond by activating the Rcs cell surface stress response pathway. LPS and divalent cations mitigate gp28-mediated growth inhibition. Additionally, gp28 inhibits growth of numerous *E. coli* strains, other Gram-negative species within the *Gammaproteobacteria*, and a representative Gram-positive bacterial strain. We propose that gp28 is attracted to negatively charged surface moieties, like LPS phosphate groups, and then disrupts bacterial membrane integrity enough to inhibit growth and eventually cause cell lysis (Fig. 4F).

### Where is Gp28 and how that affects lysis timing

Previously, we detected gp28 exclusively in membrane fractions, suggesting gp28 associates with the bacterial envelope. Here, we further show using sucrose gradient fractionation that gp28 primarily localizes to the IM prior to lysis, indicating that targeting the IM is a critical step in its lytic effect. Gp28 also permeabilizes both inner and outer membranes to small molecule dyes(12). Specific CAMPs, such as polymyxin E (colistin), with membrane-permeabilizing activity are known to dissipate IM PMF(28). Therefore, we predicted that gp28 would reduce the proton motive force leading to earlier lysis. Interestingly, gp28 accumulation in the membrane did not trigger the release of membrane-anchored SAR endolysin, but did provoke earlier holin triggering, suggesting it perturbs membrane integrity without causing a complete ablation of proton motive force. Since this earlier triggering wrests control of lysis timing from holin, it is seemingly an evolutionary paradox. How gp28 accumulation in and permeabilization of the membrane confers an advantage for phage beyond lysis requires further study.

### gp28 and CAMP membrane targets

Due to its net positive charge, we predicted gp28 would have an affinity for negatively charged membranes. The integrity of the OM depends on the electrostatic interactions between divalent cations (Mg^2+^ and Ca^2+^) and negatively charged phosphate groups in LPS, which bridging adjacent LPS molecules to stabilize and rigidify the layer (29). Externally added gp28 peptide triggered the Rcs OM stress pathway in a concentration-dependent fashion, as seen in PMB and other CAMPs (30, 31), suggesting a perturbation of the LPS barrier. The low Rcs activation from phage infection and plasmid expression of gp28 suggests that only a small amount of gp28 reaches the OM from the cytoplasm, consistent with our localization results. These findings indicate that the amount of gp28 reaching OM surface targets may differ depending on whether gp28 is expressed inside the cell or added externally.

To explore how the state of the cell surface affects gp28 activity, we explored gp28 synthetic peptide activity. Extracted LPS blocked gp28 activity in the MIC assay, consistent with an interaction that prevents gp28 entry to the cell (Fig. 2F). Further, supplementation with Mg^2+^ and Ca^2+^ in the media to reinforce the LPS leaflet reduced gp28 effectiveness by two-fold and the cation-chelator EDTA had the opposite effect (Fig. 2E), consistent with a divalent cation competitive action as seen in other CAMPs (29)(32, 33) (34). To examine the contribution of LPS charge to gp28 activity, we used LPS inner core mutants that are decorated with fewer phosphate groups. However, inner-core mutants exhibit increased OM permeability (26, 27), making their increased sensitivity to gp28 not linkable exclusively to surface charge.

Nevertheless, gp28 associates with liposomes that do not contain LPS, demonstrating a general interaction with membranes (14). Intriguingly, the Gram-positive bacterium *B. subtilis*, was susceptible to gp28-mediate growth inhibition in a teichoic acid-dependent manner (Fig. 4A). The cathelicidin LL-37 peptide demonstrates a similar activity profile (33, 35). Our current data support a model where gp28 interacts with anionic surface components such as LPS in Gram-negative bacteria and teichoic acid polymers in Gram-positive bacteria, more similar to the cathelicidins than the polymyxins (35–37). Model membrane studies revealed a higher sensitivity of the bacterial IM to low concentrations of gp28 as an intrinsic property of the bacterial membrane lipids (Fig. 4D). Therefore, future gp28 characterization can leverage the facile genetics of the phiKT phage and its host, or synthetic gp28 peptide, to elucidate the basis of its affinity for and interaction with specific lipids or lipid types.

### Expanded activity spectrum for gp28

In prior studies, we observed gp28 lethality against laboratory and environmental *E. coli* strains. Here, we expanded to a broader panel of LPS-containing Gram-negative species, demonstrating a wider activity spectrum. We also observed gp28 antimicrobial activity against a set of UPEC clinical isolates. This is consistent with data reported by Wilkinson et al. showing gp28 activity against a multidrug-resistant *P. aeruginosa* clinical isolate (38). An exception in our panel exhibiting resistance to both gp28 and PMB was *P. mirabilis*, which exhibits high intrinsic resistance to PMB and other CAMPs due to LPS modifications including lipid A palmitoylation and O-antigen acetylation that alter OM properties (39–41). We also observed that *E. coli* expressing gp28 were more susceptible to a wide range of antibiotics (Fig. 3A), consistent with a general disruption of membrane permeability.

This work supports a model in which phiKT gp28 interacts with negatively charged cell-envelope components and compromises membrane integrity, an essential part of the lysis process. Characterizing phage-encoded antimicrobial peptides that weaken the OM like this may help guide strategies to improve the entry and activity of newly developed or existing antimicrobials that cannot readily overcome the OM barrier.

## Materials and Methods

### Bacteria, phage, and growth conditions

*E. coli* and other bacterial strains, their plasmids, and phage derivatives used in this study are listed in Tables 1-4. Bacterial cultures were routinely grown in lysogeny broth (LB) at 37°C, unless otherwise noted. Lysogens and strains with toxic lysis gene clones were cultured at 30°C. All other bacterial strains used in this study were grown in Mueller-Hinton broth at 37°C and streaked on Mueller-Hinton agar plates, except *B. subtilis* strains, which were cultured in Mueller-Hinton broth and agar supplemented with 10 mM MgCl₂.

**Table 1.**
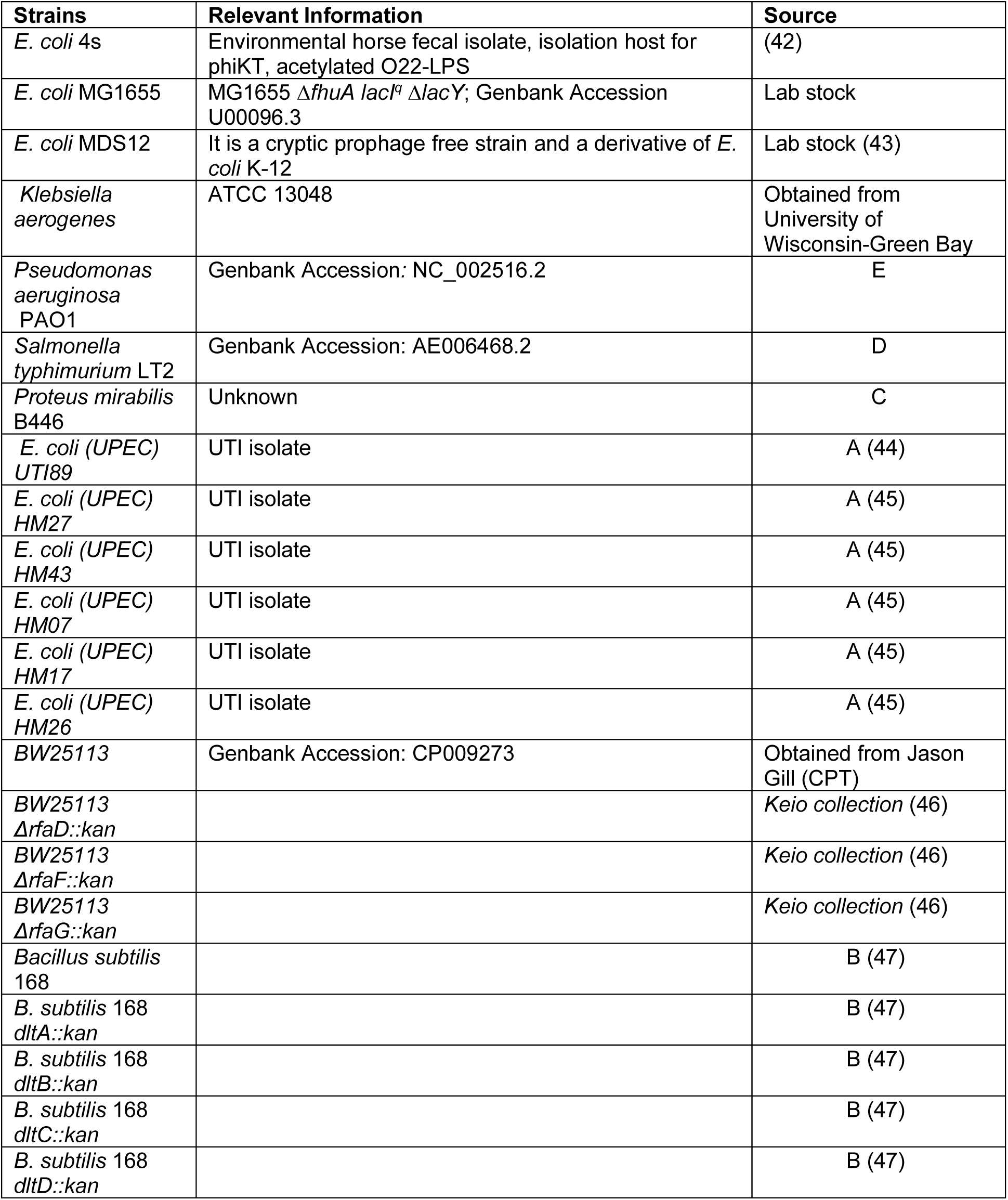
List of Bacterial strains used in this study. Sources: A-gift from Sargurunathan Subashchandrabose at Texas A&M University; B-gift from Jennifer Herman lab at Texas A&M University (BKE *B. subtilis* 168 single-gene deletion library); C-from GangaGen, gift to Center for Phage Technology; D-From Sherwood Casjens, gift to Center for Phage Technology; E-From Joe Bondy-Denomy, gift to Center for Phage Technology.

| Strains | Relevant Information | Source |
|---|---|---|
| *E. coli* 4s | Environmental horse fecal isolate, isolation host for phiKT, acetylated O22-LPS | (42) |
| *E. coli* MG1655 | MG1655 Δ*fhuA lacI*<sup>q</sup> Δ*lacY*; Genbank Accession U00096.3 | Lab stock |
| *E. coli* MDS12 | It is a cryptic prophage free strain and a derivative of *E. coli* K-12 | Lab stock (43) |
| *Klebsiella aerogenes* | ATCC 13048 | Obtained from University of Wisconsin-Green Bay |
| *Pseudomonas aeruginosa* PAO1 | Genbank Accession*:* NC_002516.2 | E |
| *Salmonella typhimurium* LT2 | Genbank Accession: AE006468.2 | D |
| *Proteus mirabilis* B446 | Unknown | C |
| *E. coli (UPEC) UTI89* | UTI isolate | A (44) |
| *E. coli (UPEC) HM27* | UTI isolate | A (45) |
| *E. coli (UPEC) HM43* | UTI isolate | A (45) |
| *E. coli (UPEC) HM07* | UTI isolate | A (45) |
| *E. coli (UPEC) HM17* | UTI isolate | A (45) |
| *E. coli (UPEC) HM26* | UTI isolate | A (45) |
| *BW25113* | Genbank Accession: CP009273 | Obtained from Jason Gill (CPT) |
| *BW25113 ΔrfaD::kan* |  | *Keio collection* (46) |
| *BW25113 ΔrfaF::kan* |  | *Keio collection* (46) |
| *BW25113 ΔrfaG::kan* |  | *Keio collection* (46) |
| *Bacillus subtilis* 168 |  | B (47) |
| *B. subtilis* 168 *dltA::kan* |  | B (47) |
| *B. subtilis* 168 *dltB::kan* |  | B (47) |
| *B. subtilis* 168 *dltC::kan* |  | B (47) |
| *B. subtilis* 168 *dltD::kan* |  | B (47) |

**Table 2.**
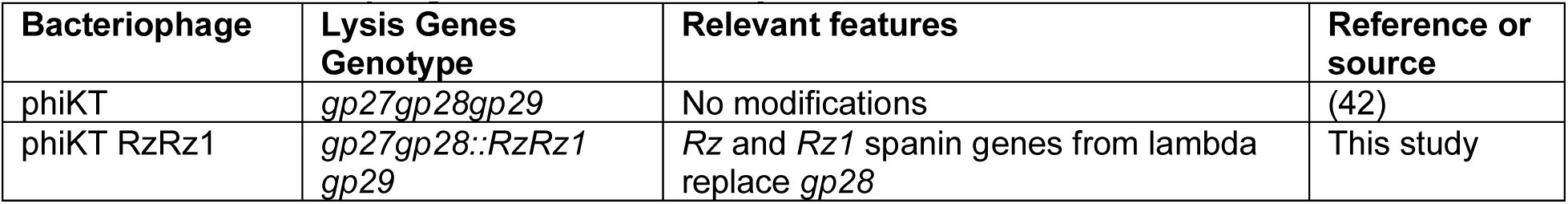
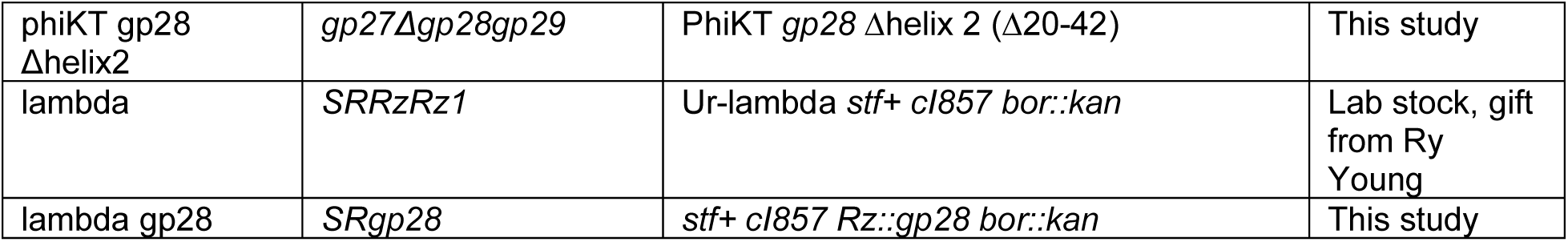
List of bacteriophages used in this study.

**Table 3.** Plasmids used in this study.

| Plasmids | Genotype and relevant features | Source |
|---|---|---|
| pBAD24 empty | pBR322-derived plasmid containing P<sub>BAD</sub> promoter, *araC* regulator, and ampicillin resistance (*bla*) | Lab stock (48) |
| pBAD24 gp28 | PhiKT gene 28 | This study |
| pBAD24 S105 | Lambda gene S105, the holin | This study |
| pBAD33 empty | pACYC184-derived plasmid containing a P<sub>BAD</sub> promoter, araC regulator, and chloramphenicol resistance (*cat*) | Lab stock (48) |
| pBAD33 gp22 | Mu gene 22, the SAR endolysin | Lab stock (49) |
| psJexD empty | pSC101-derived plasmid containing the EilR regulated PJExD promoter, the eilR repressor gene, and spectinomycin resistance | Gift from Joint BioEnergy Institute (50) |
| psJExD gp28 | PhiKT gene 28 | This study (50) |
| psJExD-spec-gp28 | PhiKT gene 28 | This study (50) |
| pQ-Kan | pQ with lambda *Q* under P*lac/ara-1* | Lab stock (51) |
| pR' Rz1 | pR' containing lambda Rz1 | Lab stock (52) |
| pAK1 gp28 | ColE1-origin, ampicillin-resistant plasmid in which the λ *P_R* late promoter drives expression of the S-R-gp28 lysis cassette genes | This study |
| pBA560-Cas13a-RzRz1 | Encodes eLbuCas13a and a spacer targeting λ RzRz1 | Gift from Jennifer Doudna, Addgene plasmid #186236 (53) |
| pJW15 empty | a *lux* reporter plasmid derived from pNLP10 and carrying a kanamycin resistance marker | Gift, Mitchell lab (54) |
| pJW15 PrprA-lux | pJW15 containing PrprA promoter transcriptionally fused to lux reporter | Gift, Mitchell lab (54) |
| pKD46 | Low copy, temperature-sensitive plasmid encoding Lambda Red recombinase genes under arabinose-inducible pBAD promoter | Lab stock (55) |

**Table 4.**
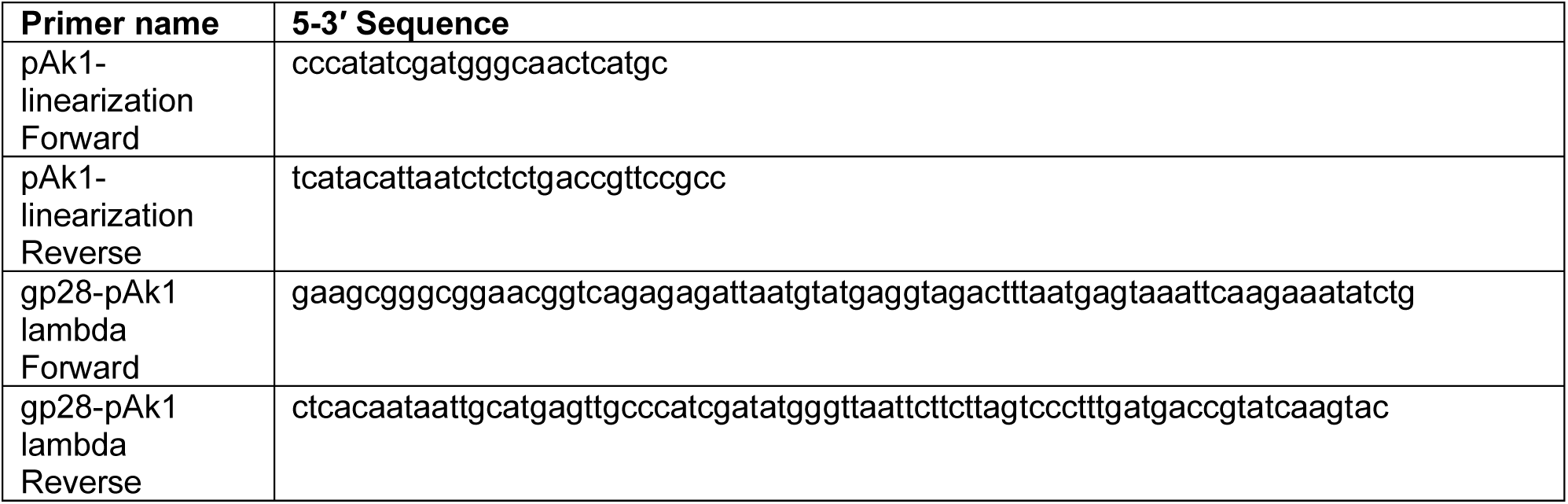
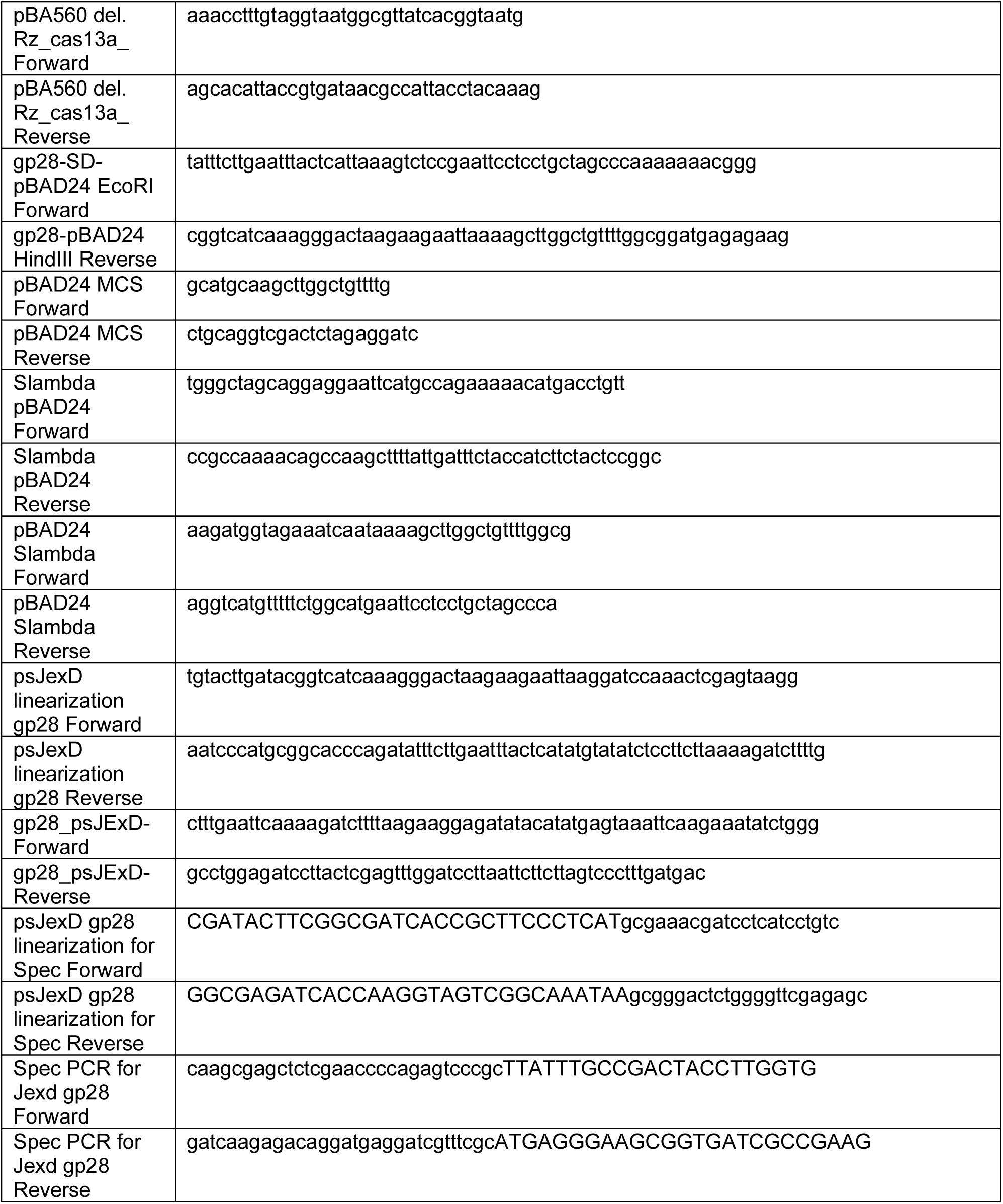
Oligonucleotides sequences used in this study.

### Bacterial growth for lysis curves

For infections and inductions, a 1:200 dilution into 25 mL of fresh LB broth was prepared from saturated overnight cultures and grown with orbital aeration to mid-log absorbance, A_550_ of ∼0.2, before induction or infection. For phage infections, mid-log cultures were infected at a multiplicity of infection (MOI) of 1, 3, or 5. For thermal induction, lysogenic strains were grown with aeration at 30°C, shifted to 42°C for 15 min to induce the prophage, and then transferred to 37°C. Chemical induction of plasmid-borne genes was conducted by addition of an inducer (isopropyl-β-D-thiogalactopyranoside (IPTG) at 1 mM, crystal violet at 25 nM, or arabinose of 1% (vol/vol)).

### Commercial peptide sources

Polymyxin B sulfate was sourced from Calbiochem (San Diego, CA). LL-37 and Nisin were purchased from Sigma-Aldrich (St. Louis, MO).

### Gp28 peptide synthesis, purification, and validation

gp28 was synthesized on a 0.1-mmol scale using Fmoc-based solid-phase peptide synthesis (SPPS), following previously reported protocols with laboratory-specific optimizations (14, 15, 56, 57). Syntheses were performed using a CEM Liberty Blue 2.0 automated peptide synthesizer equipped with a Discovery Bio microwave system. Asparagine-preloaded TGA resin (Novabiochem; loading, 0.19 mmol/g) served as the solid support, and N, N-dimethylformamide (DMF) was used as the primary solvent. Fmoc protecting groups were removed with 10% (vol/vol) piperidine in DMF. Amino acid couplings were performed under microwave-assisted conditions using 0.2 M Fmoc-protected amino acids activated with N, N′-diisopropylcarbodiimide (DIC; 15.6% [vol/vol]) and Oxyma (14.2% [wt/vol]). Following chain assembly, peptides were cleaved from the resin using a mixture of trifluoroacetic acid (TFA), triisopropylsilane (TIPS), water, and 1,2-ethanedithiol (EDT) (85:5:5:5, vol/vol/vol/vol) for at least 3 hours at room temperature. The cleavage mixture was filtered to remove the resin, and the filtrate was concentrated under a nitrogen stream. Crude peptides were precipitated with cold diethyl ether and collected by centrifugation. The resulting pellets were washed three times with ice-cold diethyl ether to remove residual cleavage reagents and byproducts (58). The crude peptides were then purified using reversed-phase high-performance liquid chromatography (RP-HPLC) using a GE HPLC system fitted with a semipreparative C4 column (Vydac 214TP; 250 × 22 mm; 10-µm particle size). Separation was performed using a gradient of solvent A (water containing 0.1% TFA) and solvent B (90% acetonitrile/10% water containing 0.1% TFA). Fractions corresponding to the target-peptide peak were collected and lyophilized (59). Peptide identity was assessed by electrospray ionization mass spectrometry (ESI-MS) (15). Purified peptides were stored long-term at –70°C and short-term at −20°C.

### Luciferase assay

The luciferase assay was performed as previously described(60) with the following modifications. Fresh overnight cultures of strains containing the pJW15-PrprA-lux plasmid were diluted and grown as described above. Each well of a clear-bottom 96-well plate was sequentially aliquoted with a total of 200 uL containing LB broth, treatment (peptides or phages), and bacterial culture to a final inoculum of A_550_ = 0.2 (∼ 4 × 10⁴ CFU). Luminescence and absorbance (A_490_) for each strain were measured every 10 minutes using a BioTek Synergy H1 plate reader (Agilent, Santa Clara, CA).

### Phage engineering and construction

The λ gp28 and phiKT RzRz1 phages were constructed according to the method of Adler *et al.*(53). Briefly, to generate recombinant phage, the host strain containing pKD46 and the homologous recombination template plasmid parent were infected at an MOI of 0.01. Cross lysates containing recombinant phage were enriched by infecting induced counterselection host strains with the cross lysate at an MOI of 0.01. Enriched lysates were plaque-purified and the parent phages were counterselected against on lawns carrying pBA560 which encodes eLbuCas13a and a spacer targeting the replaced parent gene, induced with 1 uM anhydrous tetracycline. For λ RzRz1 the spacer was generated from oligos 5’-AAACctttgtaggtaatggcgttatcacggtaatg-3’ and 5’-AGCAcattaccgtgataacgccattacctacaaag-3’ and the phiKT gp28 spacer the spacer was generated from 5’-aaacCCTTACGACCTACAAGGAGGACAGCCACACC –3’ and 5’-agcaGGTGTGGCTGTCCTCCTTGTAGGTCGTAAGG –3’. Whole genome sequences were verified at Plasmidsaurus Inc. (Louisville, KY) using Oxford Nanopore Technology long-read sequencing services.

### Membrane separation

Bacterial membrane separation was performed as described previously, except as described below(61, 62). A 1:200 dilution of saturated overnight culture was prepared in 1 L of fresh LB broth and grown with orbital aeration at 37°C to A_550_ of 0.2. Cultures were induced with 25 mM crystal violet for 15 min and centrifuged at 5,000 x g for 10 min at 4°C. The pellet was resuspended in 10 mL of buffer containing 10 mM Tris-HCl pH 7.8, 20% sucrose, and freshly prepared 1 mM phenylmethylsulfonyl fluoride. The cells were sonicated on ice (10 min total “on time” at 20% power, 15 s on / 15 s off) using a Branson SLPe sonicator (Branson Ultrasonics, Danbury, CT, USA). After clearing cell debris, the supernatant was ultracentrifuged at 185,677 x g for 2 hours at 4°C. The pellet was resuspended in 5 mL 20% sucrose and layered onto a 20/40/65% sucrose gradient for an 18-hour 4°C ultracentrifugation at 65,000 x g. A syringe aspirated approximately 500 uL of the IM and OM bands into new tubes. 5 mL of dilution buffer (50 mM Tris-HCl pH 7.8, 300 mM NaCl) was added to each tube, followed by a second 2-hour 4°C ultracentrifugation at 185,677 x g. Finally, the membrane pellets were resuspended in lysis buffer (50 mM Tris-HCl pH 7.8, 300 mM NaCl, 10% glycerol (v/v)) and stored at –70°C.

### SDS-PAGE and Western blotting

Samples were resuspended in nonreducing sample buffer and resolved on 10-20% Novex Tricine SDS-PAGE gels (Invitrogen, Thermo Fisher Scientific, Waltham, MA, USA) for 90 min at 120 V. Proteins were transferred to a 0.45-um polyvinylidene difluoride membrane (Immobilon, MilliporeSigma) using a Trans-Blot semidry transfer cell (Bio-Rad, Hercules, CA, USA) programmed for 25 V for 30 min. After blocking with 4% (wt/vol) milk in Tris-buffered saline, membranes were incubated with rabbit anti-gp28 antibody (GenScript, Piscataway, NJ, USA) or polyclonal rabbit anti-OmpA antibody (Cusabio, Catalog #: CSB-PA359226ZA01EOD, Houston, TX, USA) at a 1:2,000 dilution. IRDye 800CW goat anti-rabbit IgG secondary antibody (LI-COR Biosciences, Catalog #: 926-32211, Lincoln, NE, USA) was used at a 1:20,000 dilution for detection. Blots were scanned using an Odyssey M imaging system (LI-COR Biosciences, Lincoln, NE, USA). Densitometry was performed using Fiji ImageJ version 1.54f to measure the relative abundance of gp28 in both inner and outer membranes.

### NADH oxidation assay

The NADH oxidation assay was performed as described previously(61). Protein concentrations for each isolated membrane fraction were approximated using a NanoDrop ND-1000 spectrophotometer (Thermo Scientific). For each reaction, 100 ug of each membrane protein or input sample (pre-fractionation sonicate) was adjusted to 1 mL with 10 mM Tris-HCl buffer (pH 7.8), and 10 uL of 10 mg/mL NADH was added. Absorbance at 340 nm was measured every 30 s for 2 min for each membrane fraction and the data compiled as Absorbance (y-axis) vs Time (x-axis) for each membrane fraction.

### Minimum Inhibitory Concentration assay

Mueller-Hinton broth (not cation adjusted) was used for Minimum Inhibitory Concentration (MIC) assays, unless otherwise noted. Fresh colonies were inoculated to generate saturated overnight cultures. A 1:200 dilution from the overnight was inoculated into fresh Mueller-Hinton broth and grown to an Absorbance at 550 nm (A_550_) of ∼1.5 at 37°C. Cultures were then diluted to yield a final inoculum of ∼10^5^ CFU/mL in a total volume of 100 uL per well of a 96-well dish. Polymyxin B sulfate (Calbiochem, San Diego, CA, USA) in water, nisin from *Lactococcus lactis* (Sigma-Aldrich, St. Louis, MO, USA) in 0.05% acetic acid, chloramphenicol (Sigma-Aldrich, St. Louis, MO, USA) in ethanol, or gp28 (synthesized as previously described(15)) dissolved in water were tested at concentrations ranging from 0-512 ug/mL in triplicate. Plates were incubated for 16 hours static at 37°C. For the competition MIC assay, MH broth was first aliquoted into each well, followed by purified LPS, antibiotic or gp28, and finally the bacterial culture.

### Zone of Inhibition assay

Saturated MG1665 overnight cultures with 0.2% glucose were used at 1:40 dilution to pour a 0.75% agar lawn on LB agar plates, including 0.2%w/v% of arabinose as appropriate. The Kirby Bauer Assay used antibiotic disks discs impregnated with 30 ug of nalidixic acid, sulfamethoxazole, imipenem, ciprofloxacin, penicillin, meropenem, cefuroxime, cefoxitin, aztreonam, doxycycline, tobramycin, gentamicin, cefepime, ceftazidime, streptomycin, clarithromycin, colistin, polymyxin b, ceftriaxone, or tigecycline. After overnight incubation at 37°C, the zone of inhibition was manually measured across the disc.

### LPS extraction, purification, and detection

LPS was extracted using the hot phenol-water method as described previously(63), with modifications. A 1:200 dilution was prepared from *E. coli* 4s overnight cultures into a 1 L fresh LB broth incubated overnight at 37°C with vigorous shaking. Cultures were centrifuged at 5,000 x g for 10 min. Pellet was suspended in 10 mL SM Buffer (100 mM NaCl, 8 mM MgSO4, 50 mM Tris-HCl pH 7.5) and sonicated on ice (5 min total “on time” at 20% power, 15 s on / 15 s off) using a Branson SLPe sonicator (Branson Ultrasonics, Danbury, CT, USA). Sample was incubated for 1 hour in the presence of 1 mg DNAse I and 1 mg RNAse A. SDS was added to a final concentration of 0.5% w/v, then incubated at 65°C for 30 min. 1 mg of proteinase K was added to the sample and incubated overnight at 55°C with gentle agitation. An equivalent volume of preheated 70°C phenol was added to the sample with thorough mixing for 30 min. The mixture was cooled on ice to room temperature (RT) then centrifuged for 10 min at 5,000 x g. Two additional phenol extractions were performed on the supernatant (aqueous phase). The final aqueous phase was dialyzed against running deionized water for 2 days. The sample was then centrifuged at 100,000 x g for 1.5 hours and then precipitated overnight at –20°C with 1/10 volume of 5 M NaCl and 5 volumes of ethanol. A pellet collected from 20 min at 10,000 x g was dissolved in distilled water and centrifuged for 15 min at 15,000 x g. The ethanol precipitation step was repeated on the supernatant, which was then dialyzed to remove all traces of ethanol and further purified using a Superose 6 Increase 10/300 GL column (Cytiva) using a Bio-Rad FPLC system with 20 mM Tris-HCl pH 8.0, and 150 mM NaCl as the running buffer. LPS elution was monitored at 206 nm. Centrifugation and ethanol precipitation steps were repeated with the eluted fractions. The samples were dialyzed with two changes of water after 2 hours and left overnight. The samples were lyophilized, and the dried weight was recorded to determine the concentration of the sample. Lyophilized samples were resuspended in water and loaded on 12% Tris glycine Acrylamide/Bis-acrylamide (19:1). The gels were electrophoresed at a constant 20 mA current in Tris-glycine SDS buffer. The gel was stained using the Silver Stain Plus kit (Bio-Rad, Hercules, CA, USA) according to the manufacturer’s instructions.

### FRET-spectroscopy on liposome model membranes

Liposomes were prepared from chloroform stocks of phospholipids (L-α-phosphatidylcholine, PC; L-α-phosphatidylethanolamine from *E. coli*, PE, 840027P; L-α-phosphatidylglycerol from *E. coli*, PG, 841188P; and 1’,3’-bis [1,2-dioleoyl-*sn*-glycero-3-phospho]-glycerol, 18:1 Cardiolipin, 710335P, Avanti Polar lipids). The inner membrane (IM) model was composed of (PE:PG:CL 81:17:2 M) and the outer membrane (OM) inner leaflet model was composed of (PE:PC 70:30 M). *E.coli* Ra LPS (strain EH100, Sigma-Aldrich) was dissolved in chloroform:methanol (1:1 v/v) and prepared as an OM outer leaflet model. Lipids were mixed in the organic phase and labeled with fluorophore-conjugated NBD-PE (donor) and Rh-DHPE (acceptor) at a molar ration of 100:1:1 (M). The organic phase was dissolved under a stream of nitrogen, lipids reconstituted at 1 mM in 10 mM HEPES, 150 mM NaCl, pH 7.0 and small-unilamellar liposomes prepared by ultrasound and temperature cycling as described previously (64). Förster-resonance-energy-transfer analysis was performed as a probe-dilution assay on a Fluorolog SPEX (Jobin Yvon Inc., Edison, NJ, USA) at 37°C under constant stirring. Fluorescence intensities I_Donor_ and I_Acceptor_ were adjusted to equal intensities and recorded for 50 s until steady state of signal change upon titration of gp28. Signal ratio was calculated as I_Donor_/I _Acceptor_ and plotted against gp28 concentration.

### Graphing and statistical analyses

Data were graphed using GraphPad Prism version 11.0.1 and Fiji ImageJ version 1.54f. Raw data images and values are provided as Supplementary Information Fig. S1, Fig. S2, Fig. S3, and Dataset S1. Figure models were drawn using Linearity Curve version 6.12. Graphics for the figures were assembled in Inkscape 1.4.3 (https://inkscape.org/).

## ACKNOWLEDGEMENTS AND FUNDING SOURCES

Research reported in this publication was supported by startup funding from the Department of Biology at Texas A&M University and the National Institute of General Medical Sciences of the National Institutes of Health under award number R35GM155289 to J.R.. Additional support came from staff at the Center for Phage Technology (CPT), an Initial University Multidisciplinary Research Initiative supported by Texas A&M University and Texas AgriLife. Gary A. Lorigan would like to acknowledge support from NSF CHE–2305834 grant and Miami University. We would like to thank Angela Mitchell for strains, antibiotics, and advice, Petr Leiman for LPS purification protocols, and the Center for Phage Technology, Jennifer Herman, and Sargurunathan Subashchandrabose for bacterial strains. The authors gratefully acknowledge the excellent technical assistance from Erik Winzheim and contributions of the lab trainees Bjarne Olsson and Artur Magomedov (Division of Immunobiophysics, Research Center Borstel). A.B.S. thanks the Leibniz Science Campus InterACt, funded by the BWFGB Hamburg and the Leibniz Association (W75/2022 InterACt and “Hamburg-X Infektionsforschung”).

## REFERENCES

1. N. Mookherjee, R. E. W. Hancock, Cationic host defence peptides: Innate immune regulatory peptides as a novel approach for treating infections. Cell. Mol. Life Sci. 64, 922 (2007).

2. E. M. Kościuczuk, P. Lisowski, J. Jarczak, N. Strzałkowska, A. Jóźwik, J. Horbańczuk, J. Krzyżewski, L. Zwierzchowski, E. Bagnicka, Cathelicidins: family of antimicrobial peptides. A review. Mol. Biol. Rep. 39, 10957–10970 (2012).

3. J. Blazyk, R. Wiegand, J. Klein, J. Hammer, R. M. Epand, R. F. Epand, W. L. Maloy, U. P. Kari, A Novel Linear Amphipathic β-Sheet Cationic Antimicrobial Peptide with Enhanced Selectivity for Bacterial Lipids*. J. Biol. Chem. 276, 27899–27906 (2001).

4. M. H. A. Alaiwa, L. R. Reznikov, N. D. Gansemer, K. A. Sheets, A. R. Horswill, D. A. Stoltz, J. Zabner, M. J. Welsh, pH modulates the activity and synergism of the airway surface liquid antimicrobials β-defensin-3 and LL-37. Proc National Acad Sci 111, 18703–18708 (2014).

5. H. Jenssen, P. Hamill, R. E. W. Hancock, Peptide Antimicrobial Agents. Clin. Microbiol. Rev. 19, 491–511 (2006).

6. R. Young, Phage lysis: Three steps, three choices, one outcome. J Microbiol 52, 243–258 (2014).

7. J. S. Dewey, C. G. Savva, R. L. White, S. Vitha, A. Holzenburg, R. Young, Micron-scale holes terminate the phage infection cycle. Proc National Acad Sci USA 107, 2219–2223 (2010).

8. R. Raab, G. Neal, C. Sohaskey, J. Smith, R. Young, Dominance in lambda S mutations and evidence for translational control. J Mol Biol 199, 95–105 (1988).

9. R. Raab, G. Neal, J. Garrett, R. Grimaila, R. Fusselman, R. Young, Mutational analysis of bacteriophage lambda lysis gene S. J. Bacteriol. 167, 1035–1042 (1986).

10. J. Cahill, R. Young, “Phage Lysis: Multiple Genes for Multiple Barriers” in Ch 2 Phage Lysis: Multiple Genes for Multiple Barriers, Advances in Virus Research., (Academic Press, Elsevier, 2019), pp. 33–70.

11. M. Xu, D. K. Struck, J. Deaton, I.-N. Wang, R. Young, A signal-arrest-release sequence mediates export and control of the phage P1 endolysin. Proc Natl Acad Sci USA 101, 6415–6420 (2004).

12. A. Holt, J. Cahill, J. Ramsey, C. Martin, C. O’Leary, R. Moreland, L. T. Maddox, T. Galbadage, R. Sharan, P. Sule, J. D. Cirillo, R. Young, Phage-Encoded Cationic Antimicrobial Peptide Required for Lysis. J Bacteriol 204, e00214–21 (2021).

13. Y. A. Knirel, N. S. Prokhorov, A. S. Shashkov, O. G. Ovchinnikova, E. L. Zdorovenko, B. Liu, E. S. Kostryukova, A. K. Larin, A. K. Golomidova, A. V. Letarov, Variations in O-antigen biosynthesis and O-acetylation associated with altered phage sensitivity in Escherichia coli 4s. J Bacteriol 197, 905–912 (2015).

14. R. H. Khan, N. C. Rotich, A. Morris, T. Ahammad, B. Baral, I. D. Sahu, G. A. Lorigan, Probing the Structural Topology and Dynamic Properties of gp28 Using Continuous Wave Electron Paramagnetic Resonance Spectroscopy. J. Phys. Chem. B (2023).

15. N. C. Rotich, R. H. Khan, A. Morris, R. McCarrick, B. Baral, E. A. Okorafor, E. Faul, L. Wardrip, I. D. Sahu, G. A. Lorigan, Probing the Secondary Structure of Membrane-Bound gp28 Using Electron Spin Echo Envelope Modulation (ESEEM) Spectroscopy. J. Phys. Chem. B (2025).

16. T. J. Silhavy, D. Kahne, S. Walker, The Bacterial Cell Envelope. Cold Spring Harb. Perspect. Biol. 2, a000414 (2010).

17. H. Nikaido, Molecular basis of bacterial outer membrane permeability revisited. Microbiol. Mol. Biol. Rev. 67, 593–656 (2003).

18. C. R. H. Raetz, C. Whitfield, Lipopolysaccharide endotoxins. Biochemistry 71, 635–700 (2002).

19. Q. Li, R. Cebrián, M. Montalbán-López, H. Ren, W. Wu, O. P. Kuipers, Outer-membrane-acting peptides and lipid II-targeting antibiotics cooperatively kill Gram-negative pathogens. *Commun*. Biol. 4, 31 (2021).

20. B. Audrain, L. Ferrières, A. Zairi, G. Soubigou, C. Dobson, J.-Y. Coppée, C. Beloin, J.-M. Ghigo, Induction of the Cpx Envelope Stress Pathway Contributes to Escherichia coli Tolerance to Antimicrobial Peptides. Appl. Environ. Microbiol. 79, 7770–7779 (2013).

21. C. Maher, K. A. Hassan, The Gram-negative permeability barrier: tipping the balance of the in and the out. mBio 14, e01205–23 (2023).

22. M. Perego, P. Glaser, A. Minutello, M. A. Strauch, K. Leopold, W. Fischer, Incorporation of D-Alanine into Lipoteichoic Acid and Wall Teichoic Acid in Bacillus subtilis Identification of Genes and Regulation. J. Biol. Chem. 270, 15598–15606 (1995).

23. J. Wecke, K. Madela, W. Fischer, The absence of D-alanine from lipoteichoic acid and wall teichoic acid alters surface charge, enhances autolysis and increases susceptibility to methicillin in Bacillus subtilis. Microbiology 143, 2953–2960 (1997).

24. B. J. Schultz, E. D. Snow, S. Walker, Mechanism of d-alanine transfer to teichoic acids shows how bacteria acylate cell envelope polymers. Nat. Microbiol. 8, 1318–1329 (2023).

25. M. Cao, J. D. Helmann, The Bacillus subtilis Extracytoplasmic-Function σX Factor Regulates Modification of the Cell Envelope and Resistance to Cationic Antimicrobial Peptides. J. Bacteriol. 186, 1136–1146 (2004).

26. J. A. Yethon, E. Vinogradov, M. B. Perry, C. Whitfield, Mutation of the Lipopolysaccharide Core Glycosyltransferase Encoded by waaG Destabilizes the Outer Membrane of Escherichia coli by Interfering with Core Phosphorylation. J. Bacteriol. 182, 5620–5623 (2000).

27. A. Ebbensgaard, H. Mordhorst, F. M. Aarestrup, E. B. Hansen, The role of outer membrane proteins and lipopolysaccharides for the sensitivity of Escherichia coli to antimicrobial peptides. Front Microbiol 9, 2153 (2018).

28. Y. Wan, J. Zheng, E. W. Chan, S. Chen, Proton motive force and antibiotic tolerance in bacteria. Microb. Biotechnol. 17, e70042 (2024).

29. L. A. Clifton, M. W. A. Skoda, A. P. L. Brun, F. Ciesielski, I. Kuzmenko, S. A. Holt, J. H. Lakey, Effect of Divalent Cation Removal on the Structure of Gram-Negative Bacterial Outer Membrane Models. Langmuir 31, 404–412 (2015).

30. C. Farris, S. Sanowar, M. W. Bader, R. Pfuetzner, S. I. Miller, Antimicrobial Peptides Activate the Rcs Regulon through the Outer Membrane Lipoprotein RcsF. J. Bacteriol. 192, 4894–4903 (2010).

31. J. Meng, G. Young, J. Chen, The Rcs System in Enterobacteriaceae: Envelope Stress Responses and Virulence Regulation. Front. Microbiol. 12, 627104 (2021).

32. C.-C. H. Chen, D. S. Feingold, Locus of Divalent Cation Inhibition of the Bactericidal Action of Polymyxin B. Antimicrob. Agents Chemother. 2, 331–335 (1972).

33. K. R. Buchholz, M. Reichelt, M. C. Johnson, S. J. Robinson, P. A. Smith, S. T. Rutherford, J. G. Quinn, Potent activity of polymyxin B is associated with long-lived super-stoichiometric accumulation mediated by weak-affinity binding to lipid A. Nat. Commun. 15, 4733 (2024).

34. J. C. Stephani, L. Gerhards, B. Khairalla, I. A. Solov’yov, I. Brand, How do Antimicrobial Peptides Interact with the Outer Membrane of Gram-Negative Bacteria? Role of Lipopolysaccharides in Peptide Binding, Anchoring, and Penetration. ACS Infect. Dis. 10, 763–778 (2024).

35. A. Goode, V. Yeh, B. B. Bonev, Interactions of polymyxin B with lipopolysaccharide-containing membranes. Faraday Discuss. 232, 317–329 (2021).

36. M. M. Domingues, R. G. Inácio, J. M. Raimundo, M. Martins, M. A. R. B. Castanho, N. C. Santos, Biophysical characterization of polymyxin b interaction with LPS aggregates and membrane model systems. Pept. Sci. 98, 338–344 (2012).

37. S. Manioglu, S. M. Modaresi, N. Ritzmann, J. Thoma, S. A. Overall, A. Harms, G. Upert, A. Luther, A. B. Barnes, D. Obrecht, D. J. Müller, S. Hiller, Antibiotic polymyxin arranges lipopolysaccharide into crystalline structures to solidify the bacterial membrane. Nat. Commun. 13, 6195 (2022).

38. R. C. Wilkinson, N. E. Thomas, A. Bhatti, M. R. Burton, N. Joyce, R. E. Jenkins, Phage-Encoded Antimicrobial Peptide gp28 Demonstrates LL-37-Like Antimicrobial Activity Against Multidrug-Resistant Pseudomonas aeruginosa. PHAGE (2024).

39. A. J. McCoy, H. Liu, T. J. Falla, J. S. Gunn, Identification of Proteus mirabilisMutants with Increased Sensitivity to Antimicrobial Peptides. Antimicrob. Agents Chemother. 45, 2030–2037 (2001).

40. W.-B. Wang, I.-C. Chen, S.-S. Jiang, H.-R. Chen, C.-Y. Hsu, P.-R. Hsueh, W.-B. Hsu, S.-J. Liaw, Role of RppA in the Regulation of Polymyxin B Susceptibility, Swarming, and Virulence Factor Expression in Proteus mirabilis. Infect. Immun. 76, 2051–2062 (2008).

41. S.-S. Jiang, M.-C. Liu, L.-J. Teng, W.-B. Wang, P.-R. Hsueh, S.-J. Liaw, Proteus mirabilis pmrI, an RppA-Regulated Gene Necessary for Polymyxin B Resistance, Biofilm Formation, and Urothelial Cell Invasion. Antimicrob. Agents Chemother. 54, 1564–1571 (2010).

42. A. Golomidova, E. Kulikov, A. Isaeva, A. Manykin, A. Letarov, The Diversity of Coliphages and Coliforms in Horse Feces Reveals a Complex Pattern of Ecological Interactions. Appl. Environ. Microbiol. 73, 5975–5981 (2007).

43. V. Kolisnychenko, G. Plunkett, C. D. Herring, T. Fehér, J. Pósfai, F. R. Blattner, G. Pósfai, Engineering a Reduced Escherichia coli Genome. Genome Res 12, 640–647 (2002).

44. S. L. Chen, C.-S. Hung, J. Xu, C. S. Reigstad, V. Magrini, A. Sabo, D. Blasiar, T. Bieri, R. R. Meyer, P. Ozersky, J. R. Armstrong, R. S. Fulton, J. P. Latreille, J. Spieth, T. M. Hooton, E. R. Mardis, S. J. Hultgren, J. I. Gordon, Identification of genes subject to positive selection in uropathogenic strains of Escherichia coli: A comparative genomics approach. Proc. Natl. Acad. Sci. 103, 5977–5982 (2006).

45. A. Sintsova, A. E. Frick-Cheng, S. Smith, A. Pirani, S. Subashchandrabose, E. S. Snitkin, H. Mobley, Genetically diverse uropathogenic Escherichia coli adopt a common transcriptional program in patients with UTIs. eLife 8, e49748 (2019).

46. T. Baba, T. Ara, M. Hasegawa, Y. Takai, Y. Okumura, M. Baba, K. A. Datsenko, M. Tomita, B. L. Wanner, H. Mori, Construction of Escherichia coli K-12 in-frame, single-gene knockout mutants: the Keio collection. Mol. Syst. Biol. 2, MSB4100050 (2006).

47. B.-M. Koo, G. Kritikos, J. D. Farelli, H. Todor, K. Tong, H. Kimsey, I. Wapinski, M. Galardini, A. Cabal, J. M. Peters, A.-B. Hachmann, D. Z. Rudner, K. N. Allen, A. Typas, C. A. Gross, Construction and Analysis of Two Genome-Scale Deletion Libraries for Bacillus subtilis. Cell Syst. 4, 291–305.e7 (2017).

48. L. M. Guzman, D. Belin, M. J. Carson, J. Beckwith, Tight regulation, modulation, and high-level expression by vectors containing the arabinose PBAD promoter. J. Bacteriol. 177, 4121–4130 (1995).

49. J. S. Chamblee, J. Ramsey, Y. Chen, L. T. Maddox, C. Ross, K. H. To, J. L. Cahill, R. Young, Endolysin Regulation in Phage Mu Lysis. Mbio 13, e00813–22 (2022).

50. T. L. Ruegg, J. H. Pereira, J. C. Chen, A. DeGiovanni, P. Novichkov, V. K. Mutalik, G. P. Tomaleri, S. W. Singer, N. J. Hillson, B. A. Simmons, P. D. Adams, M. P. Thelen, Jungle Express is a versatile repressor system for tight transcriptional control. Nature communications 9, 4596–33 (2018).

51. A. Gründling, M. D. Manson, R. Young, Holins kill without warning. Proc National Acad Sci 98, 9348–9352 (2001).

52. J. Berry, E. J. Summer, D. K. Struck, R. Young, The final step in the phage infection cycle: the Rz and Rz1 lysis proteins link the inner and outer membranes. Mol Microbiol 70, 341–351 (2008).

53. B. A. Adler, T. Hessler, B. F. Cress, A. Lahiri, V. K. Mutalik, R. Barrangou, J. Banfield, J. A. Doudna, Broad-spectrum CRISPR-Cas13a enables efficient phage genome editing. Nat Microbiol 7, 1967–1979 (2022).

54. S. L. Vogt, A. D. Evans, R. L. Guest, T. L. Raivio, The Cpx Envelope Stress Response Regulates and Is Regulated by Small Noncoding RNAs. J. Bacteriol. 196, 4229–4238 (2014).

55. K. A. Datsenko, B. L. Wanner, One-step inactivation of chromosomal genes in Escherichia coli K-12 using PCR products. Proc Natl Acad Sci USA 97, 6640–6645 (2000).

56. F. Guzmán, M. Aróstica, T. Román, D. Beltrán, A. Gauna, F. Albericio, C. Cárdenas, Peptides, solid-phase synthesis and characterization: Tailor-made methodologies. Electron. J. Biotechnol. 64, 27–33 (2023).

57. S. Chandrudu, P. Simerska, I. Toth, Chemical Methods for Peptide and Protein Production. Molecules 18, 4373–4388 (2013).

58. D. L. Drew, T. Ahammad, R. A. Serafin, B. J. Butcher, K. R. Clowes, Z. Drake, I. D. Sahu, R. M. McCarrick, G. A. Lorigan, Solid phase synthesis and spectroscopic characterization of the active and inactive forms of bacteriophage S21 pinholin protein. Anal Biochem 567, 14–20 (2019).

59. C. T. Mant, Y. Chen, Z. Yan, T. V. Popa, J. M. Kovacs, J. B. Mills, B. P. Tripet, R. S. Hodges, Peptide Characterization and Application Protocols. Pept. Charact. Appl. Protoc. 386, 3–55 (2007).

60. A. K. Rai, J. F. Carr, D. E. Bautista, W. Wang, A. M. Mitchell, ElyC and Cyclic Enterobacterial Common Antigen Regulate Synthesis of Phosphoglyceride-Linked Enterobacterial Common Antigen. mBio 12, e02846–21 (2021).

61. M. B. Cian, N. P. Giordano, J. A. Mettlach, K. E. Minor, Z. D. Dalebroux, Separation of the cell envelope for Gram-negative bacteria into inner and outer membrane fractions with technical adjustments for Acinetobacter baumannii. J. Vis. Exp.: JoVE 10.3791/60517 (2020).

62. S. Shu, W. Mi, Separating Inner and Outer Membranes of Escherichia coli by EDTA-free Sucrose Gradient Centrifugation. Bio-Protoc. 13, e4638 (2023).

63. C. Galanos, O. Lüderitz, O. Westphal, A New Method for the Extraction of R Lipopolysaccharides. Eur. J. Biochem. 9, 245–249 (1969).

64. A. B. Schromm, L. Paulowski, Y. Kaconis, F. Kopp, M. Koistinen, A. Donoghue, S. Keese, C. Nehls, J. Wernecke, P. Garidel, E. Sevcsik, K. Lohner, S. Sanchez-Gomez, G. Martinez-de-Tejada, K. Brandenburg, M. Brameshuber, G. J. Schütz, J. Andrä, T. Gutsmann, Cathelicidin and PMB neutralize endotoxins by multifactorial mechanisms including LPS interaction and targeting of host cell membranes. Proc National Acad Sci 118, e2101721118 (2021).

