## Supplementary Figures for "Phage peptide promotes bacterial cell death via inner and outer membrane targets"


**Author Contributions**: Conceptualization: M.A.D., J.R.; Data curation: M.A.D., J.R.; Funding Acquisition: G.A.L, A.B.S, J.R.; Investigation: M.A.D, L.A.C, B.K.B., M.S., L.W., A.B.S, J.R.; Project administration: J.R.; Resources: G.A.L.; Visualization: M.A.D., A.B.S, J.R.; Writing-original draft: M.A.D., J.R.; Writing-review & editing: M.A.D, L.A.C, B.K.B., M.S., L.W., G.A.L., A.B.S, J.R..

**This PDF file includes:**

Figures S1 to S2

Description for Dataset S1

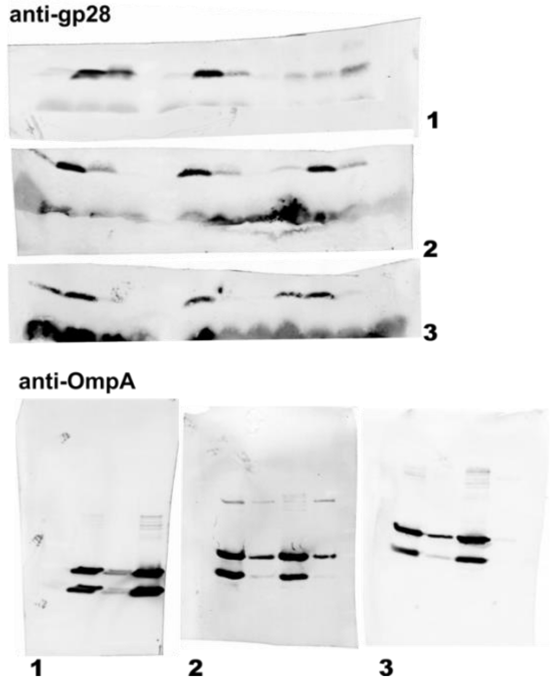


**Figure S1. Uncropped replicate blots for data shown in Fig. 1C.** Isolated membrane fractions were assayed for OM by OmpA or gp28 abundance by western blot. See main text for details.

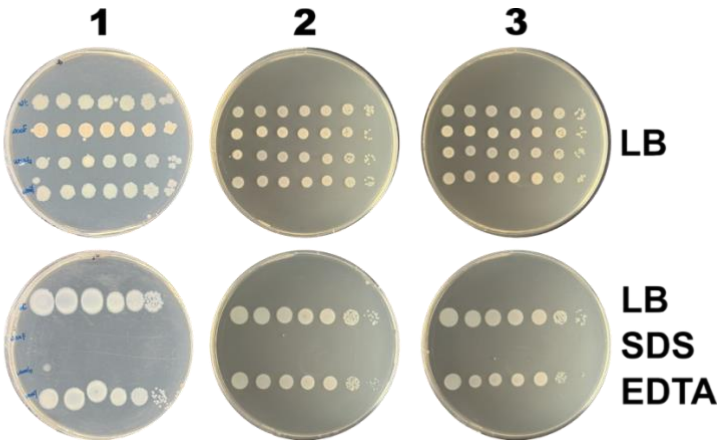


**Figure S2. Uncropped replicate plates for data shown in Fig. 4C.** Isolated membrane fractions were assayed for OM by OmpA or gp28 abundance by western blot. See main text for details.

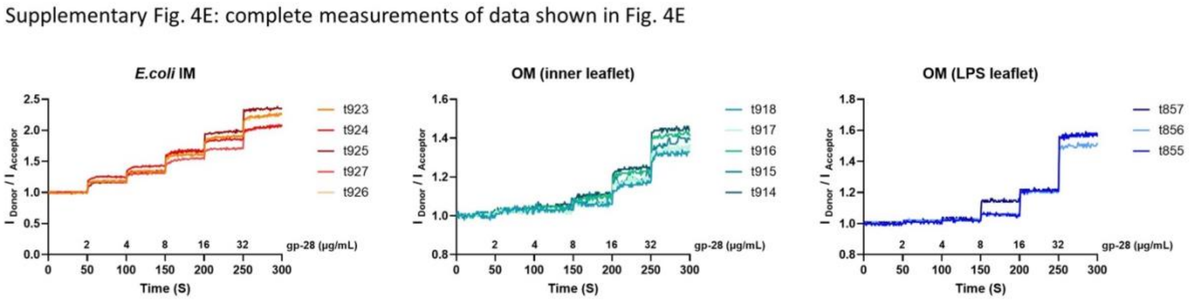


**Figure S3. Complete measurement of data shown in Fig. 4E. (A)** MG1655 Liposomal model membranes of the *E.coli* inner membrane (PE:PG:CL 81:17:2 M ), the inner leaflet of the outer membrane (PE:PC 70:30 M), and the outer leaflet of the outer membrane (*E.coli* Ra LPS) were analyzed for gp28 membrane integration by Förster-resonance-energy-transfer (FRET) probe-dilution assay. Gp28 was titrated to the liposomes at 2, 4, 8, and 32 µg/mL at 37°C to liposomes in a cuvette under constant stirring. FRET ratios ($I_{Donor}/I_{Acceptor}$) are mean ±SD of n = 5 (IM, OM inner leaflet) and n = 3 (LPS) independent measurements. See main text for details.

**Dataset S1. Raw numerical data values for graphs shown in Figures 1-4.** Provided as a separate Excel file.
